# The coronavirus proofreading exoribonuclease mediates extensive viral recombination

**DOI:** 10.1101/2020.04.23.057786

**Authors:** Jennifer Gribble, Andrea J. Pruijssers, Maria L. Agostini, Jordan Anderson-Daniels, James D. Chappell, Xiaotao Lu, Laura J. Stevens, Andrew L. Routh, Mark R. Denison

## Abstract

Coronaviruses (CoVs) emerge as zoonoses and cause severe disease in humans, demonstrated by the SARS-CoV-2 (COVID-19) pandemic. RNA recombination is required during normal CoV replication for subgenomic mRNA (sgmRNA) synthesis and generates defective viral genomes (DVGs) of unknown function. However, the determinants and patterns of CoV recombination are unknown. Here, we show that divergent β-CoVs SARS-CoV-2, MERS-CoV, and murine hepatitis virus (MHV) perform extensive RNA recombination in culture, generating similar patterns of recombination junctions and diverse populations of DVGs and sgmRNAs. We demonstrate that the CoV proofreading nonstructural protein (nsp14) 3’-to-5’ exoribonuclease (nsp14-ExoN) is required for normal CoV recombination and that its genetic inactivation causes significantly decreased frequency and altered patterns of recombination in both infected cells and released virions. Thus, nsp14-ExoN is a key determinant of both high fidelity CoV replication and recombination, and thereby represents a highly-conserved and vulnerable target for virus inhibition and attenuation.

## INTRODUCTION

Coronaviruses (CoVs) are a family of positive-sense, single-stranded RNA viruses with genomes ranging in size between 26 and 32 kb (Figure 1A and 3A). CoV recombination has been reported to be associated with increased spread, severe disease, and vaccine failure in livestock CoV epidemics (Chen et al., 2017; Feng et al., 2018) and proposed to be important in the emergence of human CoVs including the beta-CoVs (β-CoVs) severe acute respiratory syndrome coronavirus (SARS-CoV) (Anthony et al., 2017; Drosten et al., 2003; Hon et al., 2008; Ksiazek et al., 2003; Li et al., 2005) and Middle East respiratory syndrome coronavirus (MERS-CoV) (Yusof et al., 2017; Zaki et al., 2012). The ongoing severe global pandemic of SARS-CoV-2, the etiological agent of coronavirus disease 2019 (COVID-19) (Wu et al., 2020; Zhou et al., 2020) underlines the importance of defining the determinants of CoV evolution and emergence into human populations (Paraskevis et al., 2020; Randhawa et al., 2020; Wahba et al., 2020; Wong et al., 2020; Zhou et al., 2020). Studies comparing CoV strains that are closely related to SARS-CoV-2 have proposed that SARS-CoV-2 acquired the ability to infect human cells through recombination within the spike protein sequence (Huang et al., 2020; Li et al., 2020; Patiño-Galindo et al., 2020). Further, a study of genetic variation in patient SARS-CoV-2 samples has suggested that recombination may be occurring during infections in humans (Yi, 2020). Together, these data support the hypothesis that generation of novel CoVs, cross-species movement, and adaptation may be driven by recombination events in nature (Lau et al., 2015), and that CoV recombination can result in increased virulence, pathogenicity, and potential pandemic spread.

**Figure 1.**
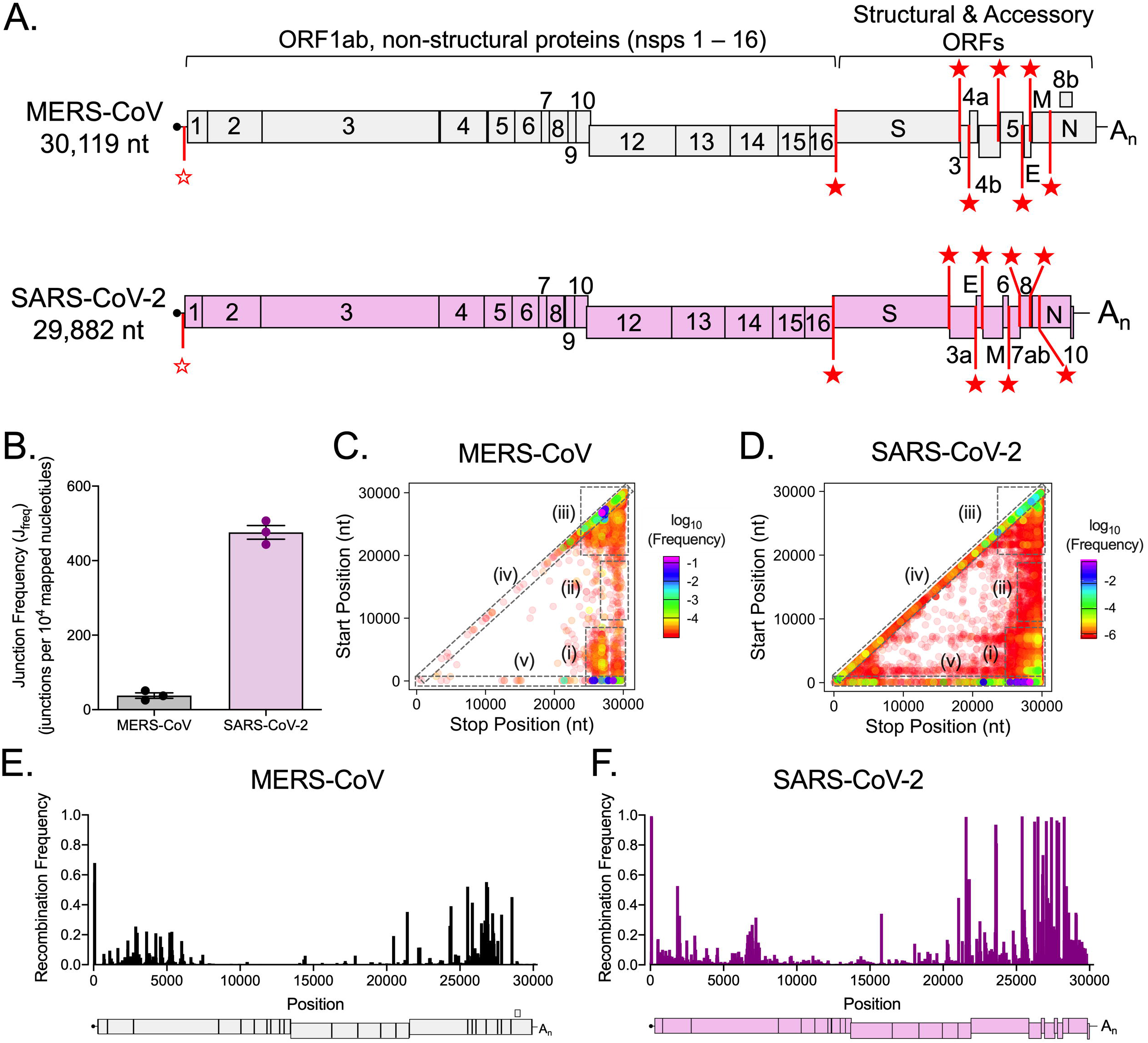
Genome-wide recombination generates populations of diverse RNA molecules in MERS-CoV and SARS-CoV-2. (A) Genome organization of MERS-CoV (violet) and SARS-CoV-2 (gray). Nonstructural (nsps 1 – 16) and structural (S, E, M, N) and accessory open reading frames (ORFs) are labelled. The common 5’ leader transcription leader sequence (TRS-L) is denoted with an unfilled red star. Body TRSs are labelled with filled red stars. MERS-CoV total cell lysates (black) and SARS-CoV-2 infected cell monolayers (violet) were sequenced by RNA-seq. (B) Junction frequency (J_freq_) was calculated by comparing the number of nucleotides in *ViReMa*-detected junctions to all mapped nucleotides. Error bars represent standard errors of the mean for three independent sequencing libraries (N = 3). Recombination junctions are mapped according to their genomic position (5’ junction site, Start Position; 3’ junction site, Stop Position) and colored according to their frequency in the population of all junctions in MERS-CoV (C) and SARS-CoV-2 (D). The highest frequency junctions are magenta and completely opaque. The lowest frequency junctions are red and the most transparent. Dashed boxes represent clusters of junctions: (i) 5’ → 3’; (ii) mid-genome → 3’ UTR; (iii) 3’ → 3’; (iv) local deletions; (v) 5’ UTR → rest of genome. Recombination frequency is quantified across the MERS-CoV (E) and SARS-CoV-2 (F) genomes. Recombination frequency is represented as a mean of three independent sequencing libraries (N = 3). See also Figure S1.

Experimental recombination has been most studied in the model β-CoV murine hepatitis virus (MHV). During mixed infections of related MHV strains, ~25% of progeny virions are generated by recombination (Keck et al., 1988; Kottier et al., 1995; Lai et al., 1985; Makino et al., 1986). During normal replication, the putative CoV replication-transcription complex (RTC) performs discontinuous transcription at virus-specific transcription regulatory sequences (TRSs) (Figure 1A, 3A) to generate a set of subgenomic mRNAs (sgmRNAs) with common 5’ leader sequences and 3’ ends, which are subsequently translated into structural and accessory proteins (Dufour et al., 2011; Kirchdoerfer and Ward, 2019; Subissi et al., 2014; Weiss et al., 1994). CoV replication also generates defective viral genomes (DVGs) that contain multiple deletions of genomic sequence. DVGs range in size from <1kb to >20kb, but retain intact 5’ and 3’ genomic untranslated regions (5’ and 3’ UTRs) and are amplified by RTC machinery supplied by co-infecting full-length helper CoVs (Brian and Spaan, 1997; Makino et al., 1985; Schaad and Baric, 1994; Wu and Brian, 2010). Therefore, CoVs perform recombination as a normal part of their replication, producing complex populations of recombined RNA molecules. Prior to the advent of next generation sequencing, direct analysis of recombined CoV RNAs was not possible and the determinants of recombination could not be identified.

In other RNA virus families including picornaviruses and alphaviruses, regulation of recombination has been mapped to replication fidelity determinants in the viral RNA-dependent RNA polymerase (RdRp) (Kempf et al., 2016; Li et al., 2019; Poirier et al., 2015; Woodman et al., 2018). In contrast to these viruses, CoV replication fidelity is primarily determined by the 3’-5’ exoribonuclease encoded in nonstructural protein 14 (nsp14-ExoN) that proofreads RNA during replication through excision of mismatched incorporated nucleotides (Ferron et al., 2018a; Ma et al., 2015). Viral exonucleases are essential for recombination in DNA viruses, including vaccinia virus (Gammon and Evans, 2009) and herpes simplex virus 1 (Grady et al., 2017; Schumacher et al., 2012). In contrast, the role of the CoV nsp14-ExoN in RNA recombination has not been defined. Catalytic inactivation of nsp14-ExoN resulted in qualitatively reduced abundance of MHV sgmRNA2 (Eckerle et al., 2007) and altered human CoV 229E (HCoV-229E) sgmRNA detection during viral recovery (Minskaia et al., 2006). These studies suggest that nsp14-ExoN RNA proofreading activity of CoVs may play a key role in RNA recombination in addition to known functions in replication fidelity, viral fitness and virulence, resistance to nucleoside analogs, and immune antagonism (Case et al., 2017; Ferron et al., 2018a; Graepel et al., 2019).

In this study, we sought to define the frequency and patterns of recombination of divergent β-CoVs SARS-CoV-2, MERS-CoV, and MHV. We show that all three viruses perform extensive recombination during replication *in vitro*, with broadly similar patterns of recombination, generating diverse populations of recombined molecules. We further demonstrate that genetically engineered inactivation of MHV nsp14-ExoN activity results in a significant decrease in recombination frequency, altered recombination junction patterns across the genome, and altered junction site selection. These defects and alterations result in a marked change in MHV-ExoN(-) recombined RNA populations, including defective viral genomes (DVGs). Previous studies our lab and others confirm the requirement of ExoN activity in high-fidelity replication, exclusion of aberrant nucleotides, viral fitness, and virulence (Eckerle et al., 2007, 2007; Ferron et al., 2018a; Graham et al., 2012). Combined with the multiple critical integrated functions of nsp14-ExoN, the demonstration in this study that nsp14-ExoN activity is required for WT-like recombination further defines it as an exceptionally conserved, vulnerable, and highly specific target for inhibition by antiviral treatments and viral attenuation. These results also support future studies aimed at illuminating the role of SARS-CoV-2 nsp14-ExoN activity in RNA recombination, the regulation of sgmRNA expression, and its contribution to novel CoV zoonotic emergence.

## RESULTS

### SARS-CoV-2 and MERS-CoV undergo extensive RNA recombination to generate populations of recombination junctions

We first sought to quantify recombination frequency and identify recombination patterns in zoonotic CoVs by sequencing both MERS-CoV and SARS-CoV-2 RNA.

Vero CCL-81 cells were infected with MERS-CoV derived from an infectious clone at passage 1 with an MOI of 0.3 pfu/mL for 72 hours until >70% cytopathic effect was observed. Vero E6 cells were infected with SARS-CoV-2 derived from a patient isolate passaged in culture (passage 5) at an MOI of 0.45 pfu/mL for 60 hours until >70% cytopathic effect was observed. Total RNA from infected cells was isolated and poly(A)-selected to capture all genomic and subgenomic RNA, as well as defective viral genomes (DVGs). Equal amounts of total cell RNA from both viruses was sequenced by short-read Illumina RNA-sequencing (RNA-seq) and long-read direct RNA Nanopore sequencing. The depth and low error rate of RNA-seq facilitated the quantification and detection of both high- and low-abundance unique junctions, but did not allow for detection of junctions in the context of a full-length RNA molecule. Long-read direct RNA sequencing on the Oxford Nanopore Technologies MinION platform was used to sequence complete RNA molecules. By combining both short- and long-read RNA sequencing, we accomplished high-confidence quantification and detection of recombination junctions as well as description of the genetic architectures of molecules formed by the junctions.

For RNA-seq, we generated libraries in triplicate from 2 μg samples of both MERS-CoV and SARS-CoV-2 infected cells. Reads were aligned to the respective viral genomes using a recombination-aware mapper, *ViReMa* (*Virus Re*combination *Ma*pper) (Routh and Johnson, 2014). *ViReMa* detects non-homologous recombination events resulting in subgenomic mRNAs (sgmRNAs) and defective viral genomes (DVGs) by identifying recombination junctions that generate a deletion greater than 5 base-pairs flanked on both sides by a 25 base-pair alignment. *ViReMa* aligned both recombined and non-recombined reads in the library and reported the total number of nucleotides aligned to the genome and all detected recombination junctions (Figure S1A-B).

MERS-CoV RNA contained an average of 19,367 detected junctions per dataset and SARS-CoV-2 libraries had an average of 56,082 detected junctions per dataset. To normalize for variations in library size between samples, recombination junction frequency (J_freq_) was calculated for MERS-CoV and SARS-CoV-2 (Figure 1B). J_freq_ refers to the number of nucleotides in all detected junctions normalized to total mapped nucleotides in a library. MERS-CoV had a mean J_freq_ of 37.80 junctions detected per 10,000 mapped nucleotides. SARS-CoV-2 had a mean J_freq_ of 475.7 junctions per 10,000 mapped nucleotides (Figure 1B). To define the patterns of these detected recombination junctions, we mapped forward (5’ → 3’) recombination junctions according to their genomic position (Figure 1C-D, Figure S1C-D). Both MERS-CoV and SARS-CoV-2 displayed clusters of junctions between the 5’ and 3’ ends of the genome, intermediate genomic positions and the 3’ end of the genome, and within the 3’ end of the genome (Figure 1B-C). SARS-CoV-2 also had many low-frequency junctions distributed across the genome, a major cluster of junctions representing local deletions and between the 5’ UTR and the rest of the genome, and horizontal clusters of low-frequency junctions between common start sites at position ~2000 and ~8000 and the rest of the genome (Figure 1D). Overall, these data demonstrate that extensive RNA recombination in both MERS-CoV and SARS-CoV-2 generates diverse populations of junctions with similar high-abundance clusters.

To determine whether high-frequency recombination occurs randomly across MERS-CoV and SARS-CoV-2 genomes, we calculated mean positional recombination frequency (Figure 1E-F) by comparing the number of nucleotides in detected junctions (both start and stop sites) at that position normalized to nucleotide depth at that position. The first and last 5000 nucleotides of both MERS-CoV and SARS-CoV-2 genomes showed clusters of recombination (Figure 1E-F). SARS-CoV-2 had many peaks of recombination frequency of greater than 0.50 (50%) across the genome (Figure 1F), suggesting that SARS-CoV-2 generates a diverse set of RNA molecules with distinct genetic architectures through recombination.

### Direct RNA Nanopore sequencing of MERS-CoV and SARS-CoV-2 defines the architecture of full-length genome, subgenomic RNAs, and defective viral genomes

We performed direct RNA Nanopore sequencing on the same RNA used for short-read RNA-seq. For both viruses, three independent sequencing experiments were performed on triplicate samples of 2 μg RNA for each virus. In order to remove prematurely truncated sequences, we bioinformatically filtered for Nanopore reads containing both genomic termini. Capture and sequencing of sgmRNAs and DVGs resulted in high sequencing coverage at both ends of the genome, while the relatively poorer capture of full-length genomes was limited due to their extreme length and resulting in depleted coverage in the middle of the genome (Figure 2A-B). We sequenced 132,493 MERS-CoV RNA molecules and 1,725,862 SARS-CoV-2 RNA molecules that had 85.6% and 82.2% identity to the parental genome, respectively (Table S1). We obtained 3 full-length direct RNA sequences of the SARS-CoV-2 genome containing over 29,850 consecutive nucleotides that aligned to the SARS-CoV-2 genome (Table S2). In MERS-CoV RNA, we detected 451 unique junction-containing full-length molecules (Figure 2A, Table S3). SARS-CoV-2 RNA contained distinct 172,191 junction-containing molecules (Figure 2B, Table S2).

**Figure 2.**
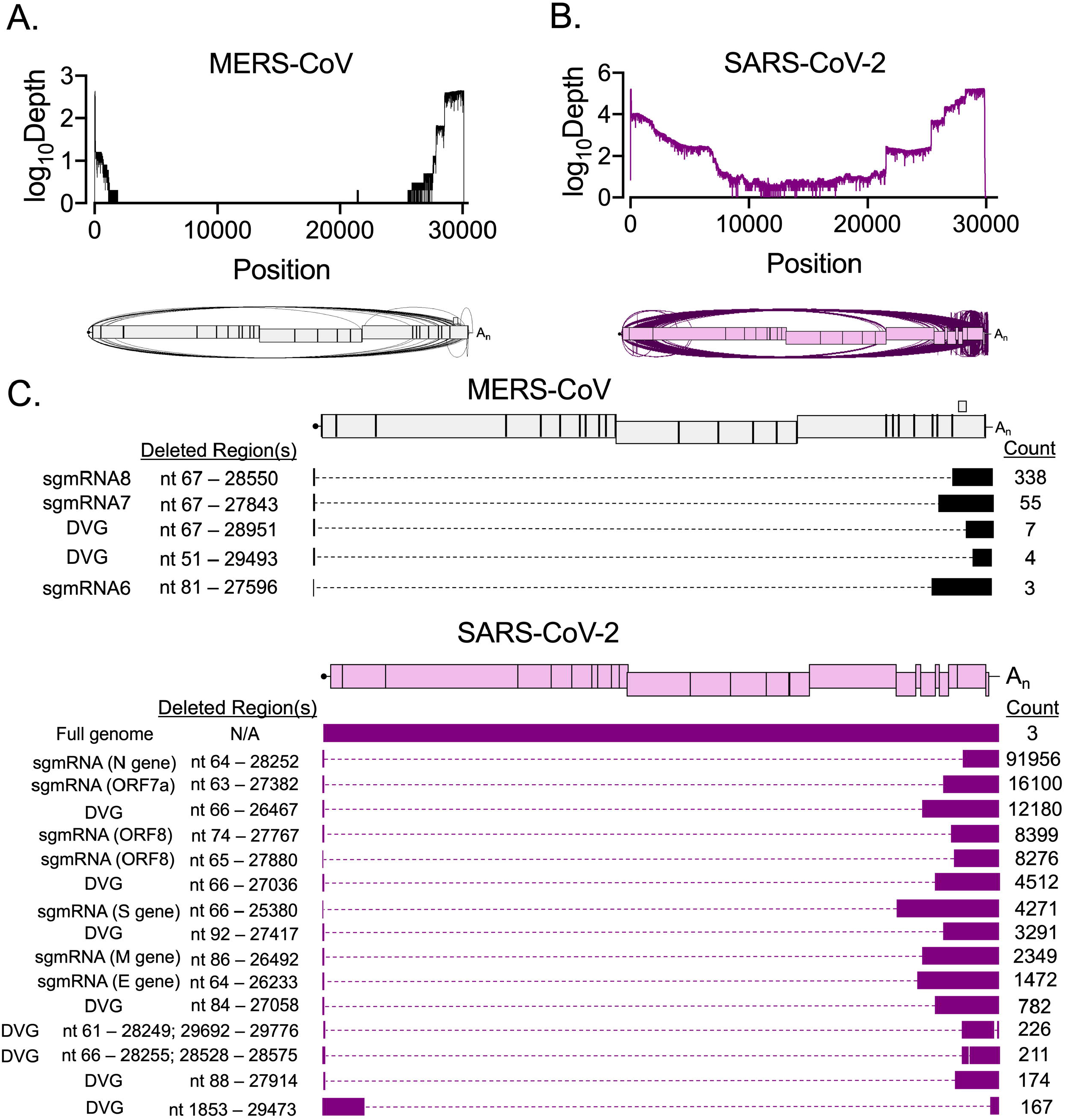
Direct RNA Nanopore sequencing of MERS-CoV and SARS-CoV-2 reveals accumulation of distinct recombined RNA populations. Direct RNA Nanopore sequencing of poly-adenylated MERS-CoV total cell lysates and SARS-CoV-2 infected cell monolayer RNA. Three sequencing experiments were performed for each virus. Nanopore reads passing quality control were combined and mapped to the viral genome using *minimap2*. Full-length reads containing the genomic termini were identified by bioinformatic filtering. Genome coverage maps and Sashimi plots visualizing junctions (arcs) in full-length (A) MERS-CoV (black) and (B) SARS-CoV-2 (violet) RNA reads. (C) Distinct RNA molecules identified in MERS-CoV (black) with at least 3 supporting reads are visualized. The number of sequenced reads containing the junction is listed (Count). Genetic sequences of each RNA molecule are represented by filled boxes and deleted regions are noted (Deleted Region(s)) and represented by dashed lines. Molecules containing junctions between the MERS-CoV TRS sequences are listed as sgmRNAs. (D) The 15 most abundant SARS-CoV-2 (violet) recombined RNA molecules and 3 full-genome reads are visualized. Molecules containing junctions linking the 5’ UTR and the predicted open reading frames of structural and accessory proteins are listed as predicted sgmRNAs. See also Table S1, Table S2, and Table S3.

To define the architectures of detected molecules, we filtered for junctions with at least 3 supporting Nanopore reads. In MERS-CoV, we defined 5 distinct species, including 3 sgmRNAs (6, 7, and 8) and 2 DVGs (Figure 2C). In SARS-CoV-2, there were 1166 isoforms with a single junction and 227 containing 2 junctions. Because the TRS sequences and sgmRNA species have not previously been experimentally confirmed in SARS-CoV-2, a Nanopore read was categorized as a predicted sgmRNA transcript if it had a junction between the predicted 5’ TRS-L and the predicted structural and accessory open reading frames to include both canonical and non-canonical sgmRNAs. The 15 most abundant isoforms in SARS-CoV-2 included 7 predicted sgmRNA transcripts and 8 DVGs (Figure 2D). We also identified potential alternative transcripts corresponding to a single gene (ORF8) (Figure 2D). In summary, direct RNA Nanopore sequencing defined a diverse set of recombined RNAs generated by both MERS-CoV and SARS-CoV-2. The results confirmed and extended the short-read RNA-seq datasets. Our results are likely an underestimate of total numbers and diversity based on the conservative bioinformatic limitation of analysis to isoforms with at least 3 supporting reads. Thus, both MERS-CoV and SARS-CoV-2 engage in extensive RNA recombination during replication in cell culture, producing diverse junctions across the viral genomes and many recombined RNA isoforms, with the indication that SARS-CoV-2 recombination exceeds that of MERS-CoV in Vero cells. These findings underline the importance of defining the determinants of CoV recombination.

### The β-coronavirus murine hepatitis virus (MHV) lacking nsp14-ExoN activity has significantly decreased and altered distribution of RNA recombination events

In picornaviruses and alphaviruses, recombination is regulated by determinants of replication fidelity (Kempf et al., 2016; Li et al., 2019; Poirier et al., 2015; Woodman et al., 2018). CoVs encode a 3’-5’ exoribonuclease (ExoN) in non-structural protein 14 (nsp14) that regulates replication fidelity by RNA proofreading (Eckerle et al., 2007; Ferron et al., 2018b; Minskaia et al., 2006). No proofreading-deficient nsp14-ExoN catalytic mutant has been rescued in MERS-CoV or SARS-CoV-2. We have rescued and extensively studied ExoN catalytic inactivation mutants (ExoN(-)) in β-CoVs murine hepatitis virus (MHV) and SARS-CoV. We used MHV and compared an engineered ExoN catalytic inactivation mutant (MHV-ExoN(-)) to wild-type virus (MHV-WT). Murine delayed brain tumor cells (DBT-9) were infected in triplicate with either MHV-WT or MHV-ExoN(-) at an MOI of 0.01 pfu/mL and total infected cell monolayer RNA was extracted when the monolayer was 95% involved in cytopathic effect and >75% intact. To isolate virions, DBT-9 cell were infected in triplicate with either MHV-WT or MHV-ExoN(-) at an MOI of 0.01 pfu/mL and virions were isolated by ultracentrifugation from supernatant collected when the monolayer was 95% involved in cytopathic effect and >90% intact. RNA-seq datasets were aligned to the MHV genome using *ViReMa*, reporting the genomic coverage in infected cells (Figure S2A-B) and virions (Figure S3A-B). To determine whether loss of nsp14-ExoN activity alters recombination, we quantified recombination junction frequency (J_freq_) in MHV-WT and MHV-ExoN(-). We found that MHV-ExoN(-) had significantly decreased J_freq_ relative to MHV-WT in both infected cells (Figure 3B) and virions (Figure 3D).

**Figure 3.**
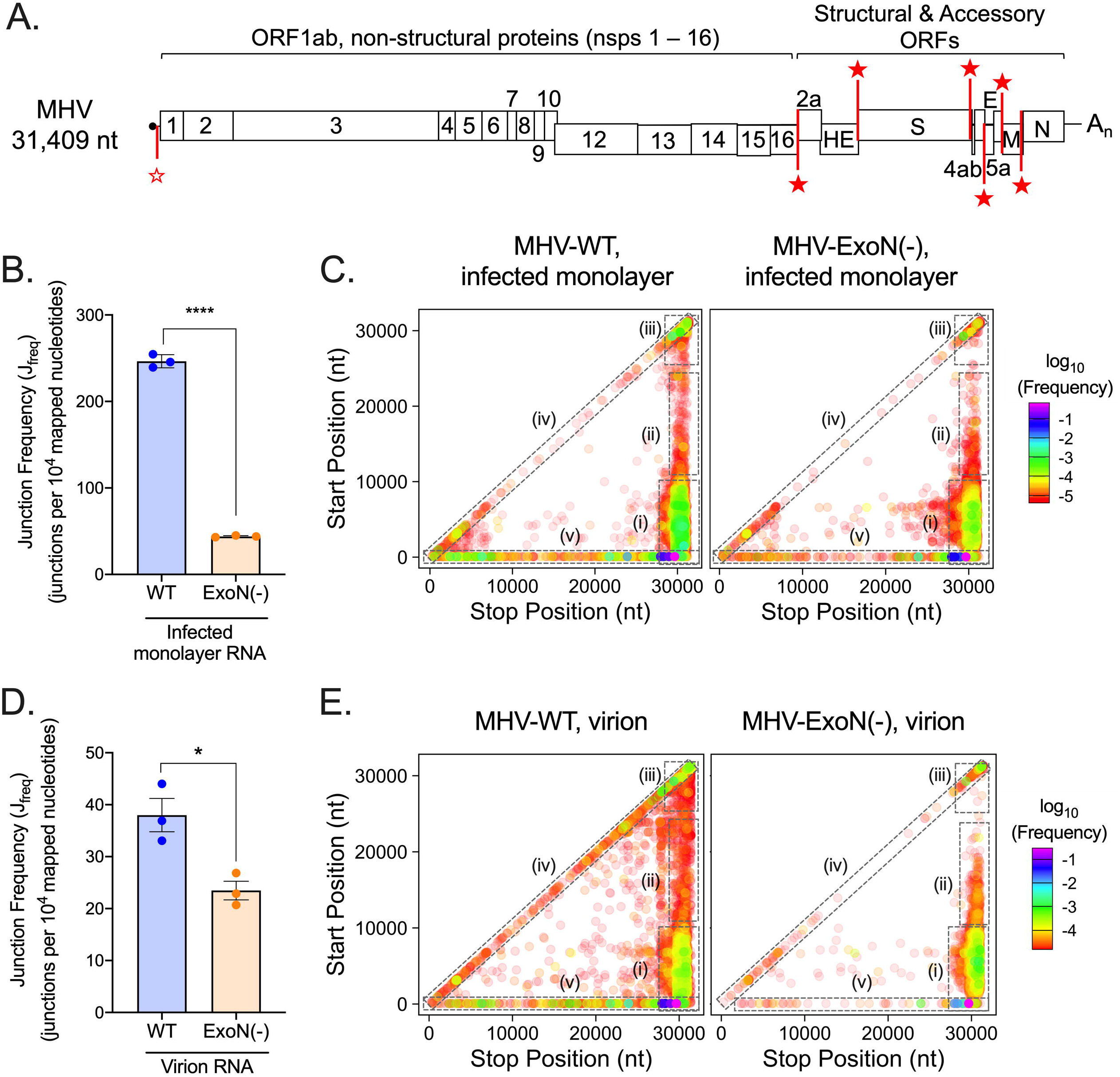
Loss of nsp14-ExoN activity decreases recombination frequency and alters recombination junction patterns across the genome. (A) Genome organization of MHV. Nonstructural (nsps 1 – 16) and structural (S, E, M, N) and accessory open reading frames (ORFs) are labelled. The common 5’ leader transcription leader sequence (TRS-L) is denoted with an unfilled red star. Body TRSs are labelled with filled red stars. Infected monolayer and virion RNA from independent experiments were poly(A) selected, sequenced by RNA-seq, and aligned to the MHV genome using *ViReMa*. Junction frequency (J_freq_) in infected monolayer RNA (B) and virion RNA (D) was calculated by dividing the number of sequenced nucleotides in all junctions by the total number of nucleotides sequenced in the library. Error bars represent standard error of the means (SEM) (N = 3). Statistical significance was determined by the unpaired student’s t-test. *, p < 0.05. Unique forward (5’ → 3’) recombination junctions detected in infected monolayers (C) and virions (E) were mapped in MHV-WT and MHV-ExoN(-) according to their genomic position. Junctions are colored according to their frequency in the population (high frequency = magenta; low frequency = red). Clusters are marked by dashed boxes: (i) 5’ → 3’; (ii) mid-genome → 3’; (iii) 3’ → 3’; (iv) local deletions; (v) 5’ UTR → rest of genome. See also Figure S2, Figure S3.

We next tested whether loss of nsp14-ExoN activity altered the population of junction sites. Recombination junctions detected by *ViReMa* were plotted according to their start (5’) and stop (3’) sites in infected cell RNA (Figure 3C, Figure S2C-D) and virion RNA (Figure 3E, Figure S3C-D). MHV-WT displayed clusters of junctions (i) between the 5’ and 3’ ends of the genome, (ii) between intermediate genomic positions and the 3’ end of the genome, (iii) between the 5’ UTR and the rest of the genome, (iv) in local deletions across the genome, and (v) within the 3’ end of the genome (Figure 3C and 3E). Similarly, MHV-ExoN(-) accumulated junction clusters between the 5’ and 3’ ends of the genome and within the 3’ end of the genome (Figure 3C and 3E). However, MHV-ExoN(-) had fewer junctions between the 5’ UTR and the rest of the genome and fewer junctions forming local deletions (Figure 3C and 3E). Together, these findings show that loss of MHV nsp14-ExoN activity resulted in decreased recombination frequency and altered junction patterns across the genome.

### Loss of nsp14-ExoN activity alters recombination in distinct genomic regions

To determine whether the engineered defect in nsp14-ExoN alters recombination across the genome, we calculated and compared mean positional recombination frequencies of MHV-WT (blue) and MHV-ExoN(-) (orange) in both infected cell monolayers (Figure 4A) and virions (Figure 4B). MHV-WT has high recombination frequency at the 5’ and 3’ ends of the genome as well as at distinct sites across the genome. MHV-ExoN(-) has a similar distribution in both infected cells and virus particles (Figure 4A-B). There were 765 positions in infected cell RNA and 499 positions in virion RNA (Table S4) with significantly altered recombination frequency in MHV-ExoN(-) compared to MHV-WT. These differences are best illustrated when the distinct genomic regions are separately visualized (Figure S4).

**Figure 4.**
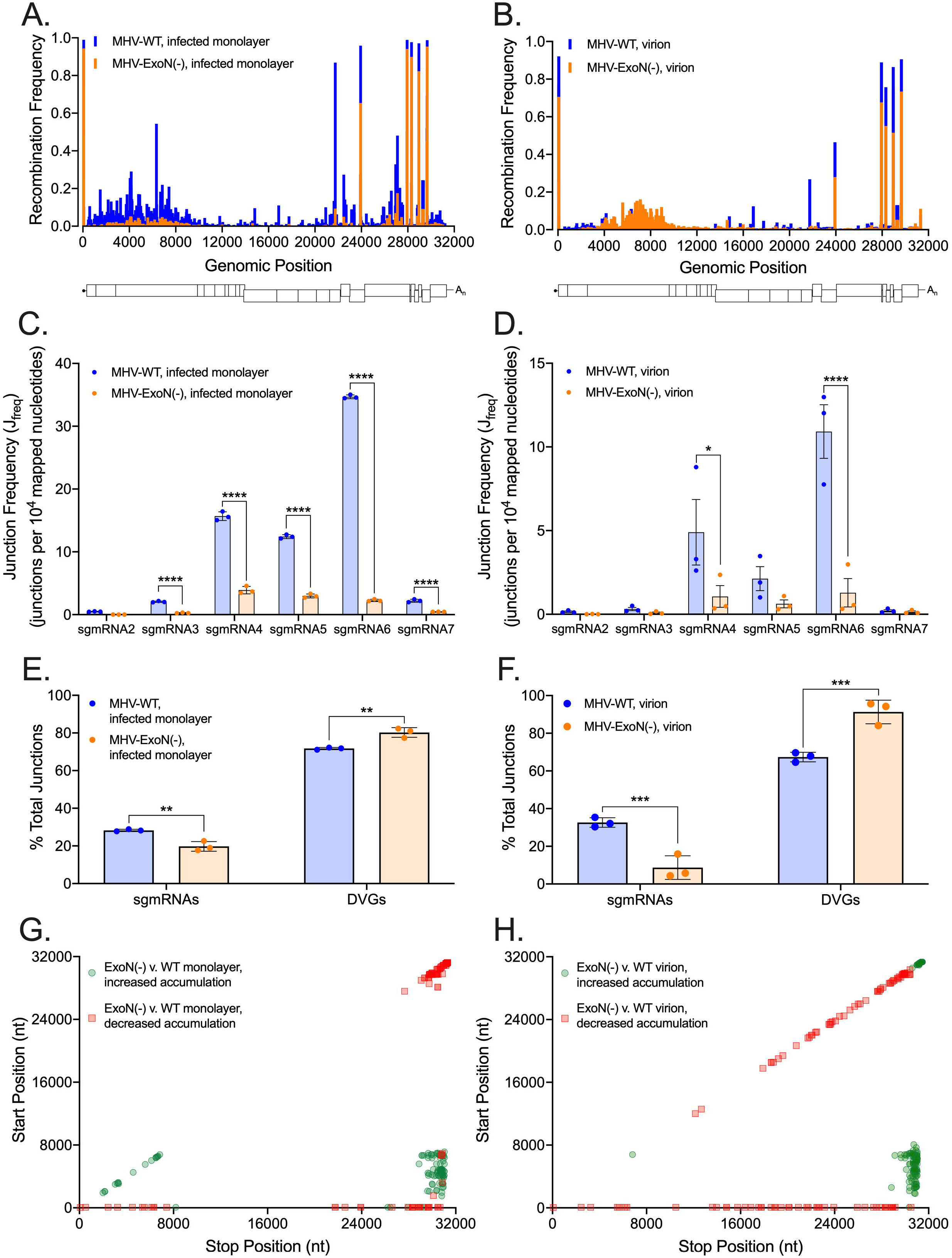
Loss of nsp14-ExoN alters recombination at multiple genomic loci and skews recombined RNA populations. Mean recombination frequency at each position across the MHV genome was compared in MHV-WT (blue) and MHV-ExoN(-) (orange) infected monolayer (A) and virion RNA (B). Positions with significantly altered recombination frequencies were identified by a 2-way ANOVA with multiple comparisons (N=3). The junction frequencies (J_freq_) of each sgmRNA were compared in MHV-WT (blue) and MHV-ExoN(-) (orange) infected monolayers (C) and virions (D). Error bars represent standard errors of the mean (SEM) (N = 3) and statistical significance was determined by a 2-way ANOVA with multiple comparisons, * p <0.05, **** p < 0.0001. (D) The percentages of sgmRNA and DVG junctions were compared in MHV-WT (blue) and MHV-ExoN(-) (orange) infected monolayers (E) and virions (F). Error bars represent SEM (N = 3). Statistical significance was determined by a 2-way ANOVA with multiple comparisons, ** p < 0.01, *** p < 0.001. The abundance of junctions in MHV-ExoN(-) was compared to MHV-WT in infected monolayers (G) and virions (H) by *DESeq2*. Junctions with statistically significant altered abundance (p < 0.05, N = 3) in MHV-ExoN(-) are mapped across the genome and colored according to their fold-change (red squares = decreased abundance, green circles = increased abundance). See also Figures S4 and S5B, Tables S4-S5.

In the 5’ UTR, nucleotides within the TRS-leader (TRS-L, yellow) participate in recombination junctions in both MHV-WT and MHV-ExoN(-) as expected, although MHV-ExoN(-) has significantly decreased recombination frequency at all sites within the TRS-L (Figure S4A, Table S4). Both MHV-WT and MHV-ExoN(-) display recombination across the nonstructural proteins, although MHV-ExoN(-) demonstrates overall decreased recombination in these regions compared with MHV-WT (Figure S4B-C, Table S4). In virions, MHV-ExoN(-) has significantly increased recombination in nsp3, nsp4, and two positions in nsp12 (Figure S4B-C, Table S4). In addition, MHV-ExoN(-) has significantly increased recombination at one position within the 3’ untranslated region (3’-UTR) (Figure S4E, Table S4).

### MHV-ExoN(-) has decreased subgenomic mRNA populations and increased abundance of defective viral genomes

We next determined whether loss of nsp14-ExoN activity influences sgmRNA patterns and abundance, as proposed in earlier studies (Eckerle et al., 2007; Minskaia et al., 2006). We quantified recombination junction frequencies (J_freq_) of each of the known junctions formed between the common TRS-L and 6 sgmRNA-specific body TRSs. sgmRNA J_freq_ refers to the number of nucleotides corresponding to the sgmRNA leader-body junction normalized to mapped library size. sgmRNAs 3, 4, 5, 6, and 7 were significantly decreased in MHV-ExoN(-) infected cell RNA (Figure 4C). sgmRNA 4 and 6 were present at significantly decreased frequencies in MHV-ExoN(-) virions compared to MHV-WT (Figure 4D). Thus, MHV-ExoN(-) has altered frequency of multiple sgmRNAs both in infected cells and in virus particles.

We further defined alterations to different populations of recombined RNA molecules in MHV-ExoN(-) compared to MHV-WT by quantifying the percentages of sgmRNA and DVG junctions in the total population of detected junctions. Any junction that did not form a canonical leader-body TRS junction was considered a DVG junction, likely including a subset of previously undefined non-canonical or alternative sgmRNA junctions. The percentage of sgmRNA junctions were significantly decreased and the percentage of DVG junctions significantly increased in both infected cell RNA (Figure 4E) and virion RNA (Figure 4F). These findings demonstrate that loss of nsp14-ExoN activity alters the locations at which the replicase machinery recombines across the genome, resulting in both decreased and altered sgmRNAs and relatively increased percentage of DVGs in MHV-ExoN(-) compared to MHV-WT. Thus, catalytic inactivation of nsp14-ExoN results in a defect in recombination site selection.

### Differential abundance of recombination junctions in MHV-ExoN(-) reveals altered recombination junction site selection

We next defined the distribution of recombination junctions with altered abundance in MHV-ExoN(-) compared to MHV-WT. We quantified the abundance junctions identified over three independent experiments and identified junction populations with altered abundances in MHV-ExoN(-) compared to MHV-WT using *DESeq2* (Love et al., 2014). MHV-ExoN(-) RNA contained distinct junction populations with both significantly increased and decreased abundance relative to MHV-WT in infected cells (Figure S5A, Table S5) and in virus particles (Figure S5B, Table S5). We mapped significantly altered junctions by their genomic position (Figure 4G-H). Recombination junctions enriched in MHV-ExoN(-) were mainly found between the 5’ and 3’ ends of the genome (Figure 4G-H). This observation suggests that MHV-ExoN(-) is accumulating altered populations of DVGs that are formed by these deletions.

Junctions with decreased abundance in MHV-ExoN(-) compared to MHV-WT in both infected cells (Figure 4G) and virions (Figures 4H) mainly clustered between the 5’ UTR and the rest of the genome. Further, infected cell RNA demonstrates a cluster of junctions with decreased abundance in MHV-ExoN(-) in the 3’ end of the genome that represent local deletions in this region (Figure 4G). This finding was complemented by the cluster of decreased abundance of local deletions of 10 – 50 bp in length across the genome in MHV-ExoN(-) virion RNA (Figure 4H). Together, these findings demonstrate that the populations of recombination junctions that were differentially abundant in MHV-ExoN(-) were not randomly distributed across the genome. This suggests that loss of nsp14-ExoN activity results in altered recombination site selection in distinct regions of the genome.

### Direct RNA Nanopore sequencing identifies defects in MHV-ExoN(-) full-length recombined RNA populations

To test the alterations of full-length recombined RNAs due to loss of nsp14-ExoN proofreading activity, we sequenced MHV-WT and MHV-ExoN(-) virion RNA by direct RNA Nanopore sequencing. Reads were mapped to the MHV genome using minimap2 and mapped reads containing both genomic termini were identified. MHV-WT virion RNA contained 101,714 complete molecules and MHV-ExoN(-) virion RNA yielded 19,334 (Table S1). Reads mapping to the full-length parental genome were not observed, likely due to the length (31.4 kb) of the MHV genome, and datasets yielded only sgmRNAs and DVGs. MHV-ExoN(-) consistently demonstrated ~1 log_10_ reduction in nucleotide depth at positions across the genome relative to MHV-WT (Figure 5A). This finding directly corroborated the decreased recombination junction frequency of MHV-ExoN(-) compared to MHV-WT demonstrated in short-read Illumina RNA-seq datasets (Figure 3B).

**Figure 5.**
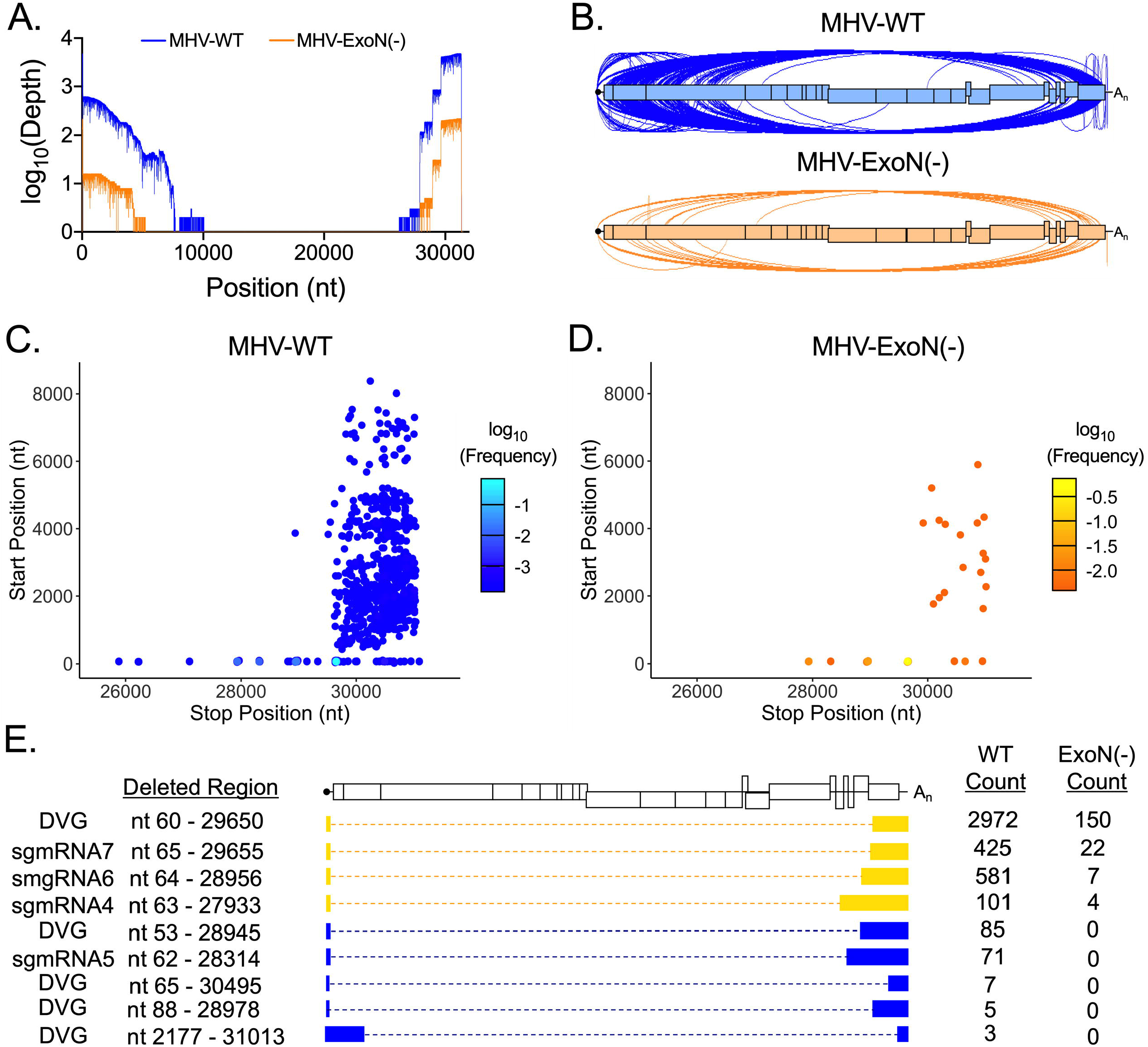
Direct RNA Nanopore sequencing of MHV full-length recombined RNA molecules. Direct RNA Nanopore sequencing of MHV virion RNA. Full-length reads were identified by bioinformatic filtering for reads containing the genomic termini. (A) Genome coverage maps of full-length MHV-WT (blue) and MHV-ExoN(-) (orange) Nanopore reads aligned to the MHV-A59 genome using *minimap2*. (B) Sashimi plot visualizing junctions (arcs) in MHV-WT (blue) and MHV-ExoN(-) (orange). (C) Junctions in reads containing only 2 discontinuous regions (1 junction) are mapped according to genome position. The junction 5’ site is mapped as the Start Position on the y-axis and the 3’ site is mapped as the Stop Position on the x-axis. Junctions are colored according to their frequency in the population (log_10_(Frequency)). MHV-WT (low = blue, high = cyan) and MHV-ExoN(-) (low = orange, high = yellow). (D) RNA molecule genetic architectures with at least 3 supporting reads identified in both MHV-WT and MHV-ExoN(-) (yellow) and unique to MHV-WT (blue). There were no reads unique to MHV-ExoN(-) supported by at least 3 reads. Genetic sequences of the RNA molecule are represented by filled boxes. Deleted regions are reported (Deleted Region) and represented by dashed lined. Canonical sgmRNA species are noted. The number of reads supporting each species are noted (Count). See also Table S1 and Table S2.

Consistent with fewer full-length molecules sequenced, MHV-ExoN(-) had a global decrease in the number of junctions across the genome (Figure 5B). Because molecules with multiple junctions in a single molecule were detected but not supported by more than one read, they were excluded from downstream analyses. We next mapped junctions detected in full-length RNA molecules according to their genomic position (Figure 5C). While MHV-WT contains 852 unique junctions present in full-length molecules, MHV-ExoN(-) had only 43 (Table S3). MHV-WT recombination junctions exclusively connect positions in the first ~8,000 nucleotides of the genome to the 3’ UTR (Figure 5C). MHV-ExoN(-) had fewer junction sites, none of which represented distinct clusters (Figure 5D). We next grouped junctions within 10-base pair windows to define distinct families of molecules supported by at least three reads. Nine such architectures were identified in MHV-WT (Figure 5E). These populations contain both DVGs and sgmRNAs. The four most abundant isoforms were also detected in MHV-ExoN(-) virion RNA, which corresponded to a DVG and sgmRNAs 4,6 and 7 (Figure 5E). We did not detect MHV-ExoN(-)-unique variants with at least 3 supporting reads, potentially due to their low frequency in the population and the limited depth of Nanopore sequencing. Additionally, all junctions generating RNA molecules found in MHV-WT and MHV-ExoN(-) were present in short-read RNA-seq datasets. These data demonstrate that loss of nsp14-ExoN activity drives the accumulation altered recombined RNA populations and DVG species diversity.

## DISCUSSION

In this study, we define the diversity of the CoV recombination landscape in MHV, MERS-CoV, and SARS-CoV-2. We also demonstrate that the loss of nsp14-ExoN activity in MHV results in decreased recombination and alters site selection of recombination junctions. Our results support a model in which nsp14-ExoN activity functions during replication to drive normal recombination, leading to the generation of populations of discrete molecules, including sgmRNAs and DVGs.

### MERS-CoV and SARS-CoV-2 have extensive recombination networks producing diverse populations of RNA species

We show that both MERS-CoV and SARS-CoV-2, the etiological agent of the ongoing global pandemic of COVID-19, performs extensive recombination and generates diverse populations of RNA molecules. SARS-CoV-2 demonstrated relatively increased recombination frequency compared to MERS-CoV and displayed unique clusters of recombination junctions across the SARS-CoV-2 genome. We analyzed SARS-CoV-2 propagated from a clinical isolate that had been passaged in cell culture, while MERS-CoV was generated from an isogenic clone of low passage. It is possible that the differences could be the result of the diversity and composition of the original sample or propagation in different cell types. Alternatively, these differences may be due to alterations in transcription and RNA synthesis kinetics in SARS-CoV-2 compared to MERS-CoV, which have not been previously defined. It will be important for future studies to determine the role of the diversity of the viral population, cell environment, and virus-specific RNA synthesis kinetics in viral recombination. Nevertheless, both viruses perform extensive, high-frequency recombination and generate diverse species of recombined RNAs.

Direct RNA Nanopore sequencing on the Oxford Nanopore Technologies MinION platform has sequenced sgmRNAs and DVGs in the alpha-coronavirus human coronavirus 229E (HCoV-229E) (Viehweger et al., 2019) and has been utilized to identify putative sgmRNAs in SARS-CoV-2 (Kim et al., 2020; Taiaroa et al., 2020). Direct RNA Nanopore sequencing further revealed the profound diversity of RNA species produced by SARS-CoV-2 during infection and yielded the first reported full-length, continuous direct RNA reads of SARS-CoV-2. The full-length genome sequences are currently the longest and most complete direct RNA Nanopore sequences reported for any CoV. Detection of multiple RNA molecules with similar but distinct genetic architectures corresponding to a single open reading frame suggests that SARS-CoV-2 produces alternative transcripts. Tandem genetic and proteomic studies of sgmRNA species in SARS-CoV-2 will be valuable to experimentally define populations of translated SARS-CoV-2 sgmRNAs their role in viral replication.

### Murine hepatitis virus (MHV) incorporates diverse recombinant RNA in its virions

Comparison of recombination in MHV-infected cells and virus particles (virions) revealed that the observed recombination patterns and species generated in infected cells are largely retained in virions. Specifically, we observed three previously unexpected recombination junction clusters: (1) junctions connecting the canonical 5’ TRS-L to non-canonical positions across the genome; (2) junctions representing small local deletions distributed evenly across the genome; and (3) local recombination within the 3’ end of the genome, including structural proteins and the 3’ UTR. This result was surprising based on earlier studies reporting the requirement of specific packaging signals for incorporation of select RNA species into the virions (Kuo and Masters, 2013). Our direct RNA Nanopore sequencing studies show that MHV virions contain a diverse population of recombined RNA molecules which exclude the canonical MHV packaging signal at position 20,273 – 20,367 (Kuo and Masters, 2013). Thus, MHV may have much lower barriers for packaging a diverse population of RNA species than previously known, which may aid in the transmission and accumulation of DVGs even under low-MOI conditions.

### MHV lacking nsp14-ExoN activity has globally decreased and altered recombination

MHV-ExoN(-) demonstrated a global decrease in recombination junction frequency in both infected cells and virions while also exhibiting skewed populations of sgmRNAs, an increased proportion of DVGs, and altered populations of recombinant isoforms. Our findings demonstrate a novel role for nsp14-ExoN in CoV RNA recombination and further suggest that loss of nsp14-ExoN activity results in altered recombination site selection. Decreased recombination frequency at canonical TRSs in MHV-ExoN(-) suggest that there is decreased accumulation of sgmRNAs. This finding is supported by previous studies (Eckerle et al., 2007; Minskaia et al., 2006) that report a defect in MHV-ExoN(-) sgmRNA synthesis. Perturbation of the abundance of each sgmRNA will likely have a profound influence on the fitness and virulence of CoVs as sgmRNAs encode critical viral proteins including nucleocapsid (N), spike (S), and many accessory ORFs that impact replication efficiency and counteract host innate immune responses (Cruz et al., 2011; Muth et al., 2018).

MHV-ExoN(-) virion RNA demonstrates a significant increase in recombination in five distinct genomic regions: 1) nsp3, 2) nsp4, 3) nsp12, 4) the spike (S) protein, and 5) the 3’ UTR. Previous studies show that CoV DVGs frequently contain sequences from the first replicase proteins after the 5’ UTR (Kim et al.; Makino; Makino et al., 1984, 1988; Penzes et al., 1996). Increased recombination in nsp3 and nsp4 may be the result of altered DVG formation and increased accumulation in virus particles. Some of the detected DVGs may be non-canonical sgmRNAs, containing junctions near the TRSs and potentially encode novel translation reading frames (Di et al., 2017; Viehweger et al., 2019).

### Loss of MHV nsp14-ExoN activity results in altered recombination site selection

Differential abundance analysis, mapping of junction detected in multiple experiments, and direct RNA Nanopore sequencing suggest that defective nsp14-ExoN activity drives altered recombination site selection resulting in different populations of accumulated molecules. In this model, abrogation of nsp14-ExoN activity would cause altered replicase activity, resulting in selection of aberrant sites, potentially based on sequence or structural elements. In both picornaviruses and alphaviruses, low fidelity mutant viruses have altered polymerase speed and processivity (Campagnola et al., 2015) and these properties contribute to recombination and the generation of defective interfering particles (Kim et al., 2019; Langsjoen et al., 2020; Poirier et al., 2015). The MHV-ExoN(-) replicase may have altered protein-protein interactions that change the stability of the complex and drive altered site selection. Alternatively, loss of nsp14-ExoN activity may result in defective replicase-RNA interactions and therefore cause altered RNA recombination site selection. In both models, nsp14-ExoN activity is an important determinant of replicase activity during RNA synthesis and recombination. However, there are likely other determinants of recombination in the CoV replicase that will be important to define through future genetic and structure-function studies.

In this study, we identify key positions across the genome that are involved in recombination junctions in the model β-CoV, MHV and we establish nsp14-ExoN proofreading activity as a key determinant of recombination in β-CoVs. Our previous studies of MHV nsp14-ExoN findings genetics and functions have been directly reproduced in SARS-CoV (Agostini et al., 2018; Eckerle et al., 2010; Smith et al., 2013). Thus, we would predict that SARS-CoV-2, will demonstrate the requirement of nsp14-ExoN activity for recombination. Further, treatment of SARS-CoV-2 infection with small molecules that target the replicase proteins including nsp14-ExoN may result in an altered recombination landscape and perturb viral fitness. It will be important for future studies to assess the role of RNA proofreading, replication fidelity, and small molecule antivirals in recombination of SARS-CoV-2. Understanding the relationship between protein determinants in RNA recombination will provide new targets for countermeasure development to combat emergence of novel CoVs from animal reservoirs with pandemic potential.

## ACKNOWLEDGMENTS

We thank members of the Denison laboratory for insightful discussions regarding this study. This work was supported by National Institutes of Health grants R01AI108197 (M.R.D.), T32GM065086 (J.G.), F31AI133952 (M.L.A). We are grateful for support from the Dolly Parton COVID-19 Research Fund, the Vanderbilt University Office of Research, and the Elizabeth B. Lamb Center for Pediatric Research at Vanderbilt University.

## AUTHOR CONTRIBUTIONS

Conceptualization (M.R.D, J.D.C., J.G.); supervision (M.R.D., A.L.R.), project administration (M.R.D., A.L.R.), funding acquisition (M.R.D., J.D.C., A.J.P.), MHV infections and deep sequencing of MHV, MERS-CoV, and SARS-CoV (J.G.), MERS-CoV infections (L.J.S.), SARS-CoV-2 infections (L.J.S), SARS-CoV-2 RNA isolation (X.L.), Nanopore sequencing (J.G.), bioinformatic analysis and visualization (J.G.), software development (J.G., A.L.R.), data curation (J.G.), writing (J.G.), review and editing (all authors).

## DECLARATION OF INTERESTS

The authors declare no competing interests.

## RESOURCE AVAILABILITY

### Lead Contact

Further information and requests for resources and reagents should be directed to and will be fulfilled by the Lead Contact, Mark R. Denison.

### Materials Availability

This study did not generate new unique reagents.

### Data and Code Availability

The datasets generated during this study are available at the Sequence Read Archive (SRA) under BioProject accession numbers PRJNA623001, PRNJA623016, PRJNA623285, PRJNA623325, PRJNA623312, PRNJA623282, PRJNA623323, PRJNA623314, PRJNA623580, PRJNA623578. The in-house scripts utilized in this study are publicly available at https://github.com/DenisonLabVU/rna-seq-pipeline.

## EXPERIMENTAL MODEL AND SUBJECT DETAILS

### Cell lines

DBT-9 (delayed brain tumor, murine astrocytoma clone 9) cells were maintained at 37°C as described previously (Chen and Baric, 1996). DBT-9 cells were originally obtained from Ralph Baric at University of North Carolina-Chapel Hill and were maintained within 50 passages of this progenitor stock. Cells were maintained in Dulbecco’s modified Eagle medium (DMEM) (Gibco) supplemented with 10% fetal clone serum (FCS) (Invitrogen), 100 U/mL penicillin and streptomycin (Gibco), and 0.25 μg/mL amphotericin B (Corning). *Cercopithecus aethiops* Vero CCL-81 cells maintained in Dulbecco’s modified Eagle medium (DMEM) (Gibco) supplemented to final concentrations of 10% fetal calf serum (Gibco), 100 IU/ml penicillin (Mediatech), 100 mg/ml streptomycin (Mediatech), and 0.25 mg/ml amphotericin B (Mediatech) were used for MERS-CoV-2 infection. Vero CCL-81 cells were obtained from ATCC. Vero E6 cells maintained in Dulbecco’s modified Eagle medium (DMEM) (Gibco) supplemented to final concentrations of 10% fetal calf serum (Gibco), 100 IU/ml penicillin (Mediatech), 100 mg/ml streptomycin (Mediatech), and 0.25 mg/ml amphotericin B (Mediatech) were used for SARS-CoV-2 infections. Vero E6 cells were obtained from ATCC.

## METHOD DETAILS

### Viruses

All MHV work was performed using the recombinant WT strain MHV-A59 (GenBank accession number AY910861.1 (Yount et al., 2002)) at passage 4 and an engineered ExoN(-) strain of MHV-A59 at passage 2. Experiments involving MERS-CoV were conducted using the human EMC/2012 strain recovered from an infectious clone (GenBank accession number JX869059.2) (Scobey et al., 2013). Experiments involving SARS-CoV-2 were conducted with a passage 5 virus inoculum generated from a Seattle, WA, USA COVID-19 patient (GenBank accession number MT020881.1). All virus manipulations were performed under stringent BSL-3 laboratory conditions according to strict protocols designed for safe and controlled handling of MERS-CoV and SARS-CoV-2.

### MHV isolation and virion purification

Subconfluent 150-cm^2^ flasks were infected with either MHV-A59 or MHV-ExoN(-) at an MOI of 0.01 PFU/cell. Supernatant was harvested at either 16 hours post infection (MHV-A59) or 24 hours post infection (MHV-ExoN(-)) when the monolayer was >95% fused and remained intact. Infection supernatant was clarified by centrifugation at 1500 × *g* for 5 minutes at 4°C. Viral particles were purified on a 30% sucrose cushion by ultracentrifugation at 25,000 RPM at 4°C for 16 hours. The viral pellet was resuspended in MSE buffer (10mM MOPS, pH 6.8; 150mM NaCl; 1 mM EDTA). Viral RNA was extracted using the TRIzol-LS reagent according to manufacturer’s protocols. RNA was quantified using the Qubit RNA HS assay. Virion data in this paper is the result of three independent experiments sequenced independently.

### MHV isolation from infected monolayers

Three subconfluent 150-cm^2^ flasks of DBT-9 cells were infected with either MHV-WT or MHV-ExoN(-) at an MOI or 0.01 PFU/cell. Monolayer was harvested at either 16 hpi (MHV-WT) or 24 hpi (MHV-ExoN(-)) when the monolayer was >95% fused and >75% intact. RNA was extracted with TRIzol according to manufacturer’s protocols. Infected monolayer data in this paper is the result of three independent experiments sequenced independently.

### MERS-CoV infection

Three nearly confluent 25-cm^2^ flasks of Vero CCL-81 cells were infected with MERS-CoV at an MOI of 0.3 pfu/cell. Total infected cell lysates were collected at 72 hpi with the monolayer < 50% intact. RNA was extracted in TRIzol according to manufacturer’s protocols.

### SARS-CoV-2 infection

Five subconfluent 25-cm^2^ flasks of Vero E6 cells were infected at an MOI = 0.45 pfu/cell and cellular monolayers were harvested 60 hpi when the monolayer was >90% fused. RNA was extracted in TRIzol according to manufacturer’s protocols.

### Short-read Illumina RNA-sequencing of viral RNA

Next generation sequencing (NGS) libraries were generated using 2 μg of RNA. RNA was submitted to Genewiz for library preparation and sequencing. Briefly, after quality control, polyadenylated RNA was selected during library preparation. Isolated RNA was heat fragmented and libraries were prepared for 2 × 250 nucleotide paired-end sequencing performed (Illumina). Genewiz performed basecalling and read demultiplexing.

### Direct RNA Nanopore sequencing

RNA from ultracentrifuge-purified virions was prepared for direct RNA Nanopore sequencing on the Oxford Nanopore Technologies MinION platform according to the manufacturer’s protocols. Libraries were sequenced on fresh MinION R9.4 flow-cells for 24 hours, or until the pore occupancy was under 20%. Virion RNA from three independent experiments was sequenced on three separate flow cells for both MHV-WT and MHV-ExoN(-). MERS-CoV RNA from three independent cultures was sequenced on three separate flow cells. SARS-CoV-2 RNA isolated from three independent infections was sequenced on three separate flow cells.

## QUANTIFICATION AND STATISTICAL ANALYSIS

### Illumina RNA-seq processing and alignment

Raw reads were processed by first removing the Illumina TruSeq adapter using *`Trimmomatic`* (Bolger et al., 2014) default settings (command line parameters java -jar trimmomatic.jar PE sample_R1.fastq.gz sample_R2.fastq.gz output_paired_R1.fastq output_unpaired_R1.fastq output_paired_R2.fastq output_unpaired_R2_unpaired.fastq ILLUMINACLIP:TruSeq3-PE.fa:2:30:10 LEADING:3 TRAILING:3 SLIDINGIWINDOW:4:15 MINLEN:36). Reads shorter than 36 bp were removed and low-quality bases (Q score < 30) were trimmed from read ends. The raw `FASTQ` files were aligned to the MHV-A59 genome (AY910861.1), the MERS-CoV genome (JX869059.2), and the SARS-CoV-2 genome (MT020881.1) using the Python2 script *`ViReMa`* (<u>V</u>iral <u>Re</u>combination <u>Ma</u>pper) (Routh and Johnson, 2014) using the command line parameters `python2 ViReMa.py reference_index input.fastq output.sam--OuputDir sample_virema/--OutputTag sample_virema-BED --MicroIndelLength 5`. The sequence alignment map (`SAM`) file was processed using the `samtools` (Li et al., 2009) suite to calculate nucleotide depth at each position in a sorted binary alignment map (`BAM`) file (using command line parameters `samtools depth -a -m θ sample_virema.sorted.bam > sample_virema.coverage`).

### Recombination junction analysis

Recombination junction frequency (J_freq_) was calculated by comparing the number of nucleotides in detected recombination junctions to the total number of mapped nucleotides in a library. Nucleotides in detected recombination junctions were quantified as a sum of nucleotide depth reported at each junction in the `BED` file generated by *`ViReMa`*. Total nucleotides mapped to the MHV-A59 genome were quantified as a sum of nucleotide depth at each position across the genome in the tab-delineated text file generated by the *`samtools`*. J_freq_ was reported as junctions per 10^4^ nucleotides sequenced. Mean J_freq_ values were compared between MHV-WT and MHV-ExoN(-) and statistical significance determined by an unpaired student’s t-test. Junctions were mapped across the genome according to their start (5’) and stop (3’) positions. These junctions were first filtered in the forward (5’ → 3’) direction using the *`dpylr`* package (RStudio). The frequency of each junction was calculated by comparing the depth of the unique junction to the total number of nucleotides in all detected junctions in a library. Junctions were plotted according to the genomic position and colored according to log_10_ of the frequency using *`ggplot2`* in RStudio.

Recombination frequency was calculated at each genomic position by dividing the number of nucleotides in any junction mapping to the position divided by the total number of nucleotides sequenced at the position. Mean recombination frequencies were compared between MHV-WT and MHV-ExoN(-) for three independent sequencing experiments by a two-way ANOVA statistical analysis with multiple comparisons.

### Identification of sgmRNA and DVG junctions

Forward recombination junctions were classified as either canonical sgmRNA junctions or DVG junctions based on the position of their junction sites and filtered in Microsoft Excel. Briefly, junction start sites were filtered to those positioned between nucleotide 62 – 72 in the genome. The stop sites were then filtered for those positioned within each respective sgmRNA TRS. sgmRNA frequency was calculated by dividing the sum of the depth of all junctions corresponding to an individual sgmRNA by the total number of mapped nucleotides at each position.

The filtered sgmRNA junctions were compiled and DVG junctions were filtered in RStudio by performing an exclusionary `anti_join()` using *`dplyr`*. DVG junctions were filtered in the forward direction. The percentage of sgmRNA and DVG junctions was calculated by comparing the depth of all filtered sgmRNA or DVG junctions to the sum of all detected forward junctions. Mean percent sgmRNA and DVG was compared between three independent sequencing experiments in virion RNA. Statistical significance was determined by a 2-way ANOVA test with multiple comparisons.

### Differential abundance of junctions

To compare the abundance of junctions in MHV-A59 and MHV-ExoN(-), the ViReMa output list of junctions was analyzed by in-house scripts (https://github.com/DenisonLabVU) and the R package *`DESeq2`* (Love et al., 2014). Junctions significantly up- or down-regulated in MHV-ExoN(-) were visualized using *`bioinfokit`* (Bedre, 2020) and further mapped according to their genomic positions. Statistical significance was determined by the p-value of each junction calculated by the `DESeq2` package in RStudio and junctions with a significant alteration of abundance in MHV-ExoN(-) compared to MHV-WT were visualized as either red or green in the graph generated by *`bioinfokit`*.

### Direct RNA Nanopore alignment and analysis

Live basecalling was performed by *`Guppy`* in *`MinKNOW`*. Run statistics were generated from each sequencing experiment by *`NanoPlot`* (De Coster et al., 2018). Pass reads from all three experiments were concatenated for each virus and aligned to the genome using `minimap2` (Li, 2018) and *`FLAIR`* (<u>F</u>ull <u>L</u>ength <u>A</u>lternative <u>I</u>soforms of <u>R</u>NA) (Tang et al., 2018) to generate alignment files and `BED` files listing deletions detected in each sequenced RNA molecule. Both `BAM` and `BED` files were filtered for full length molecules using *`samtools`* and Microsoft Excel, respectively. Full-length MHV molecules were defined as encoding coverage at positions 71 and 31034, representing the canonical TRS-L and beginning of the 3’ UTR required for replication. Because no TRS sequences have been experimentally confirmed in either MERS-CoV or SARS-CoV-2, we filtered both datasets for those that included any sequence of the 5’ UTR and 3’ UTR of the respective viruses. BED files generated by the flair align module were parsed based on the number of junctions were identified. Nanopore reads containing only 1 junction were identified using Microsoft Excel and unique junctions were quantified in RStudio using base-R functions. Sequencing coverage maps were generated from *`samtools`* depth analysis of filtered `BAM` files. All junctions present in sequenced libraries were mapped in Sashimi plots generated by the Integrated Genome Viewer (IGV) (Robinson et al., 2011). Junctions present in full-length MHV RNA molecules with a single deletion were mapped according to their genomic positions as previously described. The genetic architectures of full-length RNA molecules sequenced by direct RNA Nanopore sequencing were visualized by filtering RNA molecules for at least 3 supporting reads. Low frequency variants were removed from this analysis.

---

**Table S3. Direct RNA Nanopore read isoforms, related to Figures 2 and 5**. Direct RNA Nanopore reads aligning to viral genome by minimap2. Individual reads are listed by read name. Genomic positions of read alignment are listed (“Read Start”, “Read Stop”). Read segments aligning to the genome are noted (“# Segments”) and start positions and aligned segment legnths listed (“Segment Start”, “Segment Length”).

**Table S4. Genomic positions with significantly altered positional recombination frequency in MHV-ExoN(-) infected monolayer and virion RNA compared to MHV-WT, related to Figure 4**. Positions with significantly altered recombination frequency in MHV-ExoN(-) infected monolayer RNA compared to MHV-WT and in MHV-ExoN(-) virion RNA compared to MHV-WT as determined by a 2-way ANOVA with multiple comparisons are listed. Genomic regions are noted. (N = 3 for each infected cell and virion RNA samples)

**Table S5. Differential abundance of recombination junctions in MHV-ExoN(-) infected monolayer compared to MHV-WT, related to Figure 4**. Junctions with altered abundance in MHV-ExoN(-) infected monolayer RNA compared to MHV-WT and in MHV-ExoN(-) virion RNA compared to MHV-WT are listed. P-values calculated by DESeq2. (N = 3 for each infected monolayer and virion RNA samples)

## Notes

### Competing Interest Statement

The authors have declared no competing interest.

