## Supplemental Figures and Tables for "The coronavirus proofreading exoribonuclease mediates extensive viral recombination"

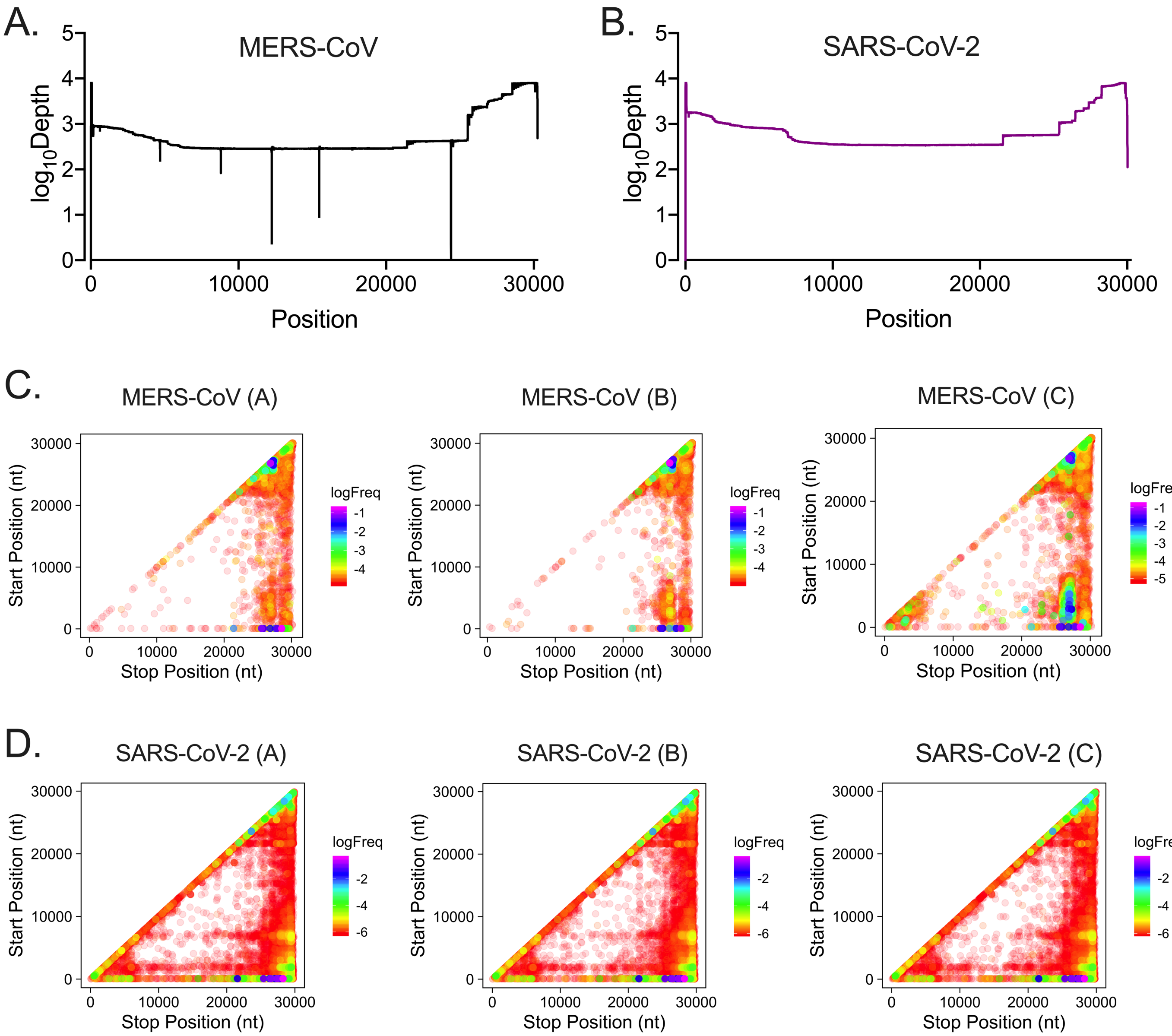


**Figure S1. Short-read RNA-sequencing genome coverage and ViReMa-detected recombination junctions in MERS-CoV and SARS-CoV-2, related to Figure 1.** RNA-seq libraries of (A) MERS-CoV and (B) SARS-CoV-2 were aligned to the viral genomes with ViReMa. Nucleotide depth was calculated at each position and represented as mean nucleotide depth (N=3). Individual recombination junction scatter plots of (C) MERS-CoV and (D) SARS-CoV-2. Recombination junctions were detected by ViReMa and forward (5' → 3') junctions were identified by bioinformatic filtering. Junctions are plotted according to their 5' (start) and 3' (stop) positions and colored according to their frequency in the population of total junctions. Highly abundant junctions are magenta and opaque and low-frequency junctions are red and transparent.

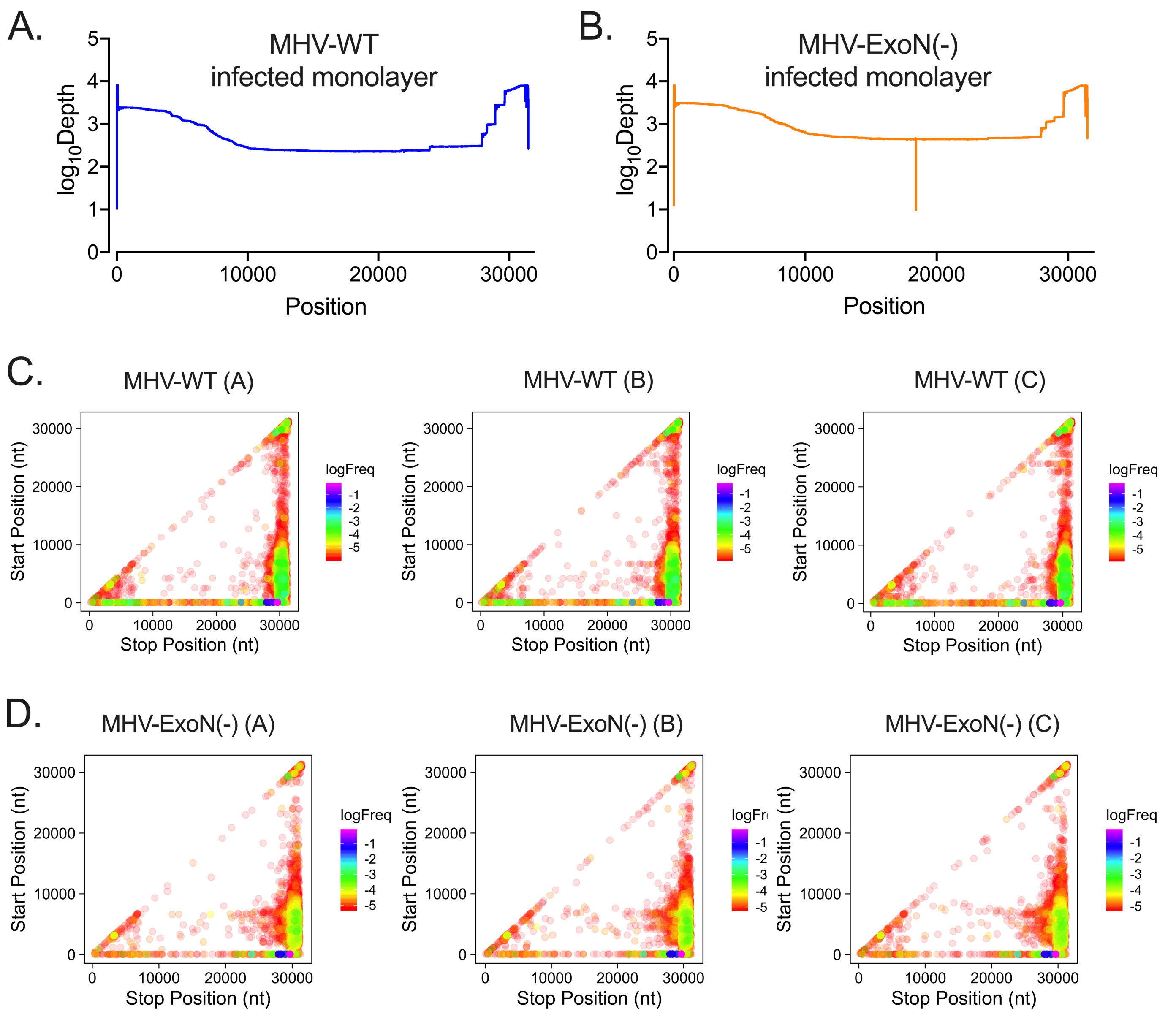


**Figure S2. Short-read RNA-sequencing genome coverage and recombination junctions detected by ViReMa in MHV monolayer RNA, related to Figure 3.** RNA-seq libraries of (A) MHV-WT and (B) MHV-ExoN(-) infected cell monolayer RNA were aligned to the viral genomes with ViReMa. Nucleotide depth was calculated at each position and represented as mean nucleotide depth (N=3). Individual recombination junction scatter plots of (C) MHV-WT and (D) MHV-ExoN(-). Recombination junctions were detected by ViReMa and forward (5' → 3') junctions were identified by bioinformatic filtering. Junctions are plotted according to their 5' (start) and 3' (stop) positions and colored according to their frequency in the population of total junctions. Highly abundant junctions are magenta and opaque and low-frequency junctions are red and transparent.

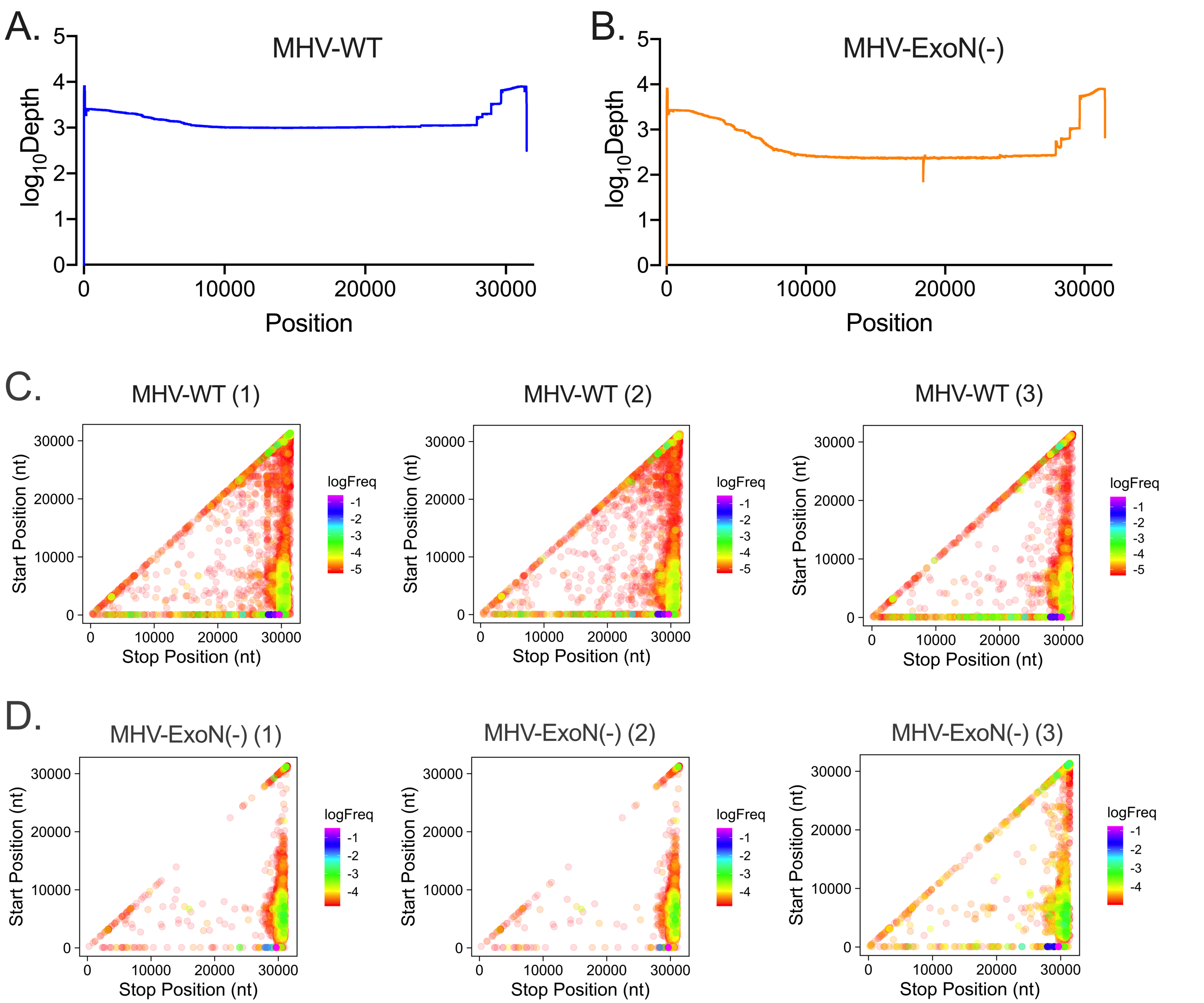


**Figure S3. Short-read RNA-sequencing genome coverage and recombination junctions detected by ViReMa in MHV virion RNA, related to Figure 3.** RNA-seq libraries of (A) MHV-WT and (B) MHV-ExoN(-) virion RNA were aligned to the viral genomes with ViReMa. Nucleotide depth was calculated at each position and represented as mean nucleotide depth (N=3). Individual recombination junction scatter plots of (C) MHV-WT and (D) MHV-ExoN(-). Recombination junctions were detected by ViReMa and forward (5' → 3') junctions were identified by bioinformatic filtering. Junctions are plotted according to their 5' (start) and 3' (stop) positions and colored according to their frequency in the population of total junctions. Highly abundant junctions are magenta and opaque and low-frequency junctions are red and transparent.

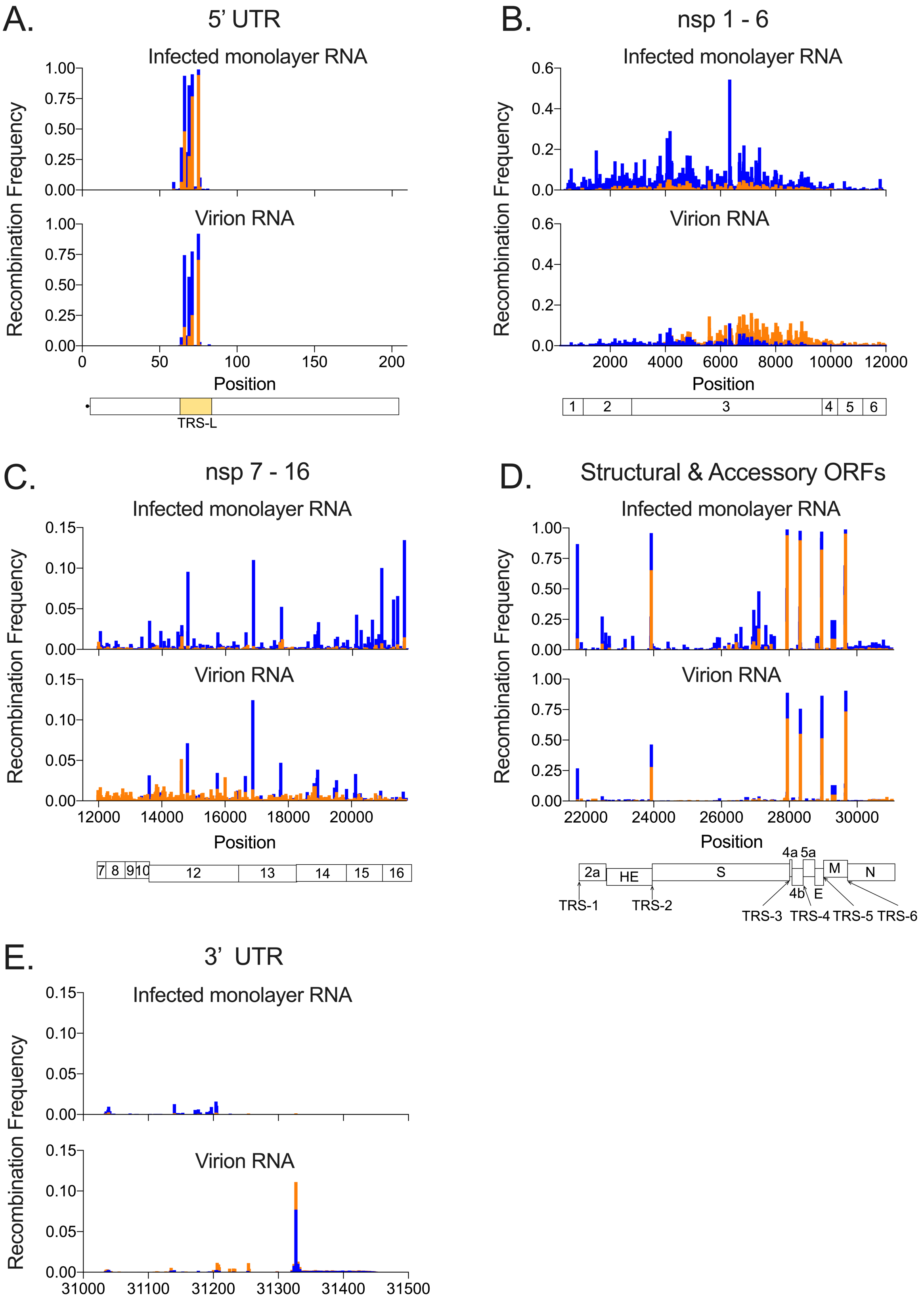


**Figure S4. MHV-ExoN(-) has significantly altered recombination frequency at multiple positions across the genome, related to Figure 4**. Mean recombination frequency at each genomic position is shown for MHV-WT (blue) and MHV-ExoN(-) (orange). (A) 5' UTR, (B) the non-replicase nonstructural proteins (nsp1 – 6), (C) the replicase proteins (nsp7 – 16), (D) the structural and accessory proteins, (E) 3' UTR. Key RNA elements including the TRS-leader (TRS-L) and body TRSs (TRS1 – 7) are labelled. Positions with statistically significant differences in MHV-ExoN(-) recombination frequency were identified by a 2-way ANOVA with multiple comparisons.

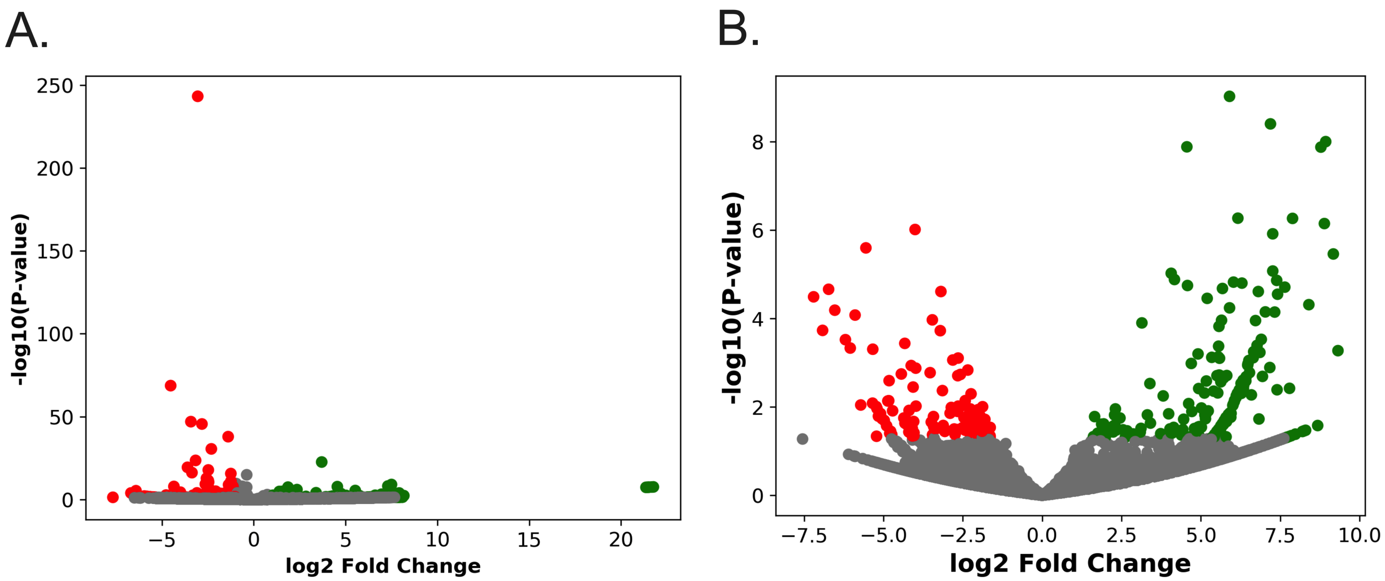


**Figure S5. MHV-ExoN(-) differentially accumulates recombination junctions in virion RNA and infected monolayers, related to Figure 4.** Recombination junction abundance was compared in MHV-ExoN(-) to MHV-WT by DESeq2 in infected cell monolayer RNA (A) and virion RNA (B). Volcano plots of ExoN(-) junctions in virions (A) and infected cell monolayers (B) colored by statistical significance (red or green, $p < 0.05$) and by the $\log_2$(Fold Change) of expression (red = downregulated, green = upregulated).

**Table S1. Alignment statistics of Nanopore direct RNA sequencing of MERS-CoV, SARS-CoV-2, MHV-WT and MHV-ExoN(-), related to Figures 2 and 5.**

| Virus | Mean % Identity | Mean Read Length | Mean Read Quality | Median % Identity | Median Read Length | Median Read Quality | Number of Reads | Read Length N50 | Total Bases |
|---|---|---|---|---|---|---|---|---|---|
| MERS-CoV | 85.6 | 773.8 | 8.4 | 85.9 | 583 | 8.4 | 132493 | 1014 | 102518766 |
| SARS-CoV-2 | 82.2 | 1555.8 | 8.9 | 83.2 | 1409 | 8.9 | 1725862 | 1952 | 2685130864 |
| MHV-WT | 86.7 | 1175.7 | 9.0 | 87.3 | 718 | 9.1 | 101714 | 1678 | 111895953 |
| MHV-ExoN(-) | 86.8 | 1062.3 | 9.1 | 87.5 | 672 | 9.2 | 19334 | 1483 | 20159852 |

**Table S2. Full genome reads of SARS-CoV-2 detected by direct RNA Nanopore sequencing, related to Figure 2.**

| Genome | Read Start (nt) | Read End (nt) | Read Name | Read Length | Count |
|---|---|---|---|---|---|
| MT02088.1 | 10 | 29691 | 103efdf4-a528-46e3-b5bb-b360e2cae18b;0 | 29681 | 1 |
| MT02088.1 | 11 | 29863 | 41da8a52-cb9e-4969-95eb-5fd13b65584b;0 | 29852 | 1 |
| MT02088.1 | 14 | 29874 | cb66c733-0ad3-493c-8a9f-310bbd96e6fe;0 | 29860 | 1 |
